# A tale of two paths: The evolution of mitochondrial recombination in bivalves with doubly uniparental inheritance

**DOI:** 10.1101/2022.10.22.513339

**Authors:** Chase H. Smith, Brendan J. Pinto, Mark Kirkpatrick, David M. Hillis, John M. Pfeiffer, Justin C. Havird

## Abstract

In most animals, mitochondrial DNA is strictly maternally inherited and non-recombining. One exception to these assumptions is called doubly uniparental inheritance (DUI): a phenomenon involving the independent transmission of female and male mitochondrial genomes. DUI is known only from the molluscan class Bivalvia. The phylogenetic distribution of male mitochondrial DNA in bivalves is consistent with several evolutionary scenarios, including multiple independent gains, losses, and varying degrees of recombination with female mitochondrial DNA. In this study, we use phylogenetic methods to test male mitochondrial DNA origination hypotheses and infer the prevalence of mitochondrial recombination in bivalves with DUI. Phylogenetic modeling using site concordance factors supported a single origin of male mitochondrial DNA in bivalves coupled with recombination acting over long evolutionary timescales. Ongoing mitochondrial recombination is present in Mytilida and Venerida, which results in a pattern of concerted evolution of female and male mitochondrial DNA. Mitochondrial recombination could be favored to offset the deleterious effects of asexual inheritance and maintain mitonuclear compatibility across tissues. Cardiida and Unionida have gone without recent recombination, possibly due to an extension of the *COX2* gene in male mitochondrial DNA. The loss of recombination may be neutral but could be connected to the role of M mtDNA in sex determination or sexual development. Our results support recombination events in DUI species may occur throughout their genomes. Future investigations may reveal more complex patterns of inheritance of recombinants, which could explain the retention of signal for a single origination of male mitochondrial DNA in protein coding genes.

## Introduction

Mitochondria are found in almost all eukaryotic cells and possess their own independently inherited mitochondrial DNA (mtDNA). Typically, animal mtDNA is ~16 kb long and contains 37 genes (13 protein-coding, two rRNAs, and 22 tRNAs) and a control region (a non-coding region that often contains the origin of replication) (Boore 1999). In most bilaterian animals, mtDNA is assumed to be strictly maternally inherited and non-recombining. However, exceptions to these generalizations have been documented across multiple phyla (Piganeau, Gardner and Eyre-Walker 2004; Barr, Neiman and Taylor 2005; Tsaousis *et al*. 2005; Ghiselli *et al*. 2021). One such exception occurs in molluscan bivalves, where several lineages show doubly uniparental inheritance (DUI). This unusual mode of mitochondrial inheritance is characterized by the transmission of two mitochondrial genomes, one passed by females to all offspring and a second passed by males to only male offspring (Hoeh, Blakley and Brown 1991; Skibinski, Gallagher and Beynon 1994). Females only possess F-mtDNA, while males are globally heteroplasmic in their somatic tissues and exclusively possess M mtDNA in their sperm (Breton *et al*. 2017, 2022; Ghiselli *et al*. 2019; Bettinazzi *et al*. 2020).

Doubly uniparental inheritance has been described from five bivalve orders: Cardiida, Mytilida, Nuculanida, Unionida, and Venerida (Gusman *et al*. 2016; Capt *et al*. 2020). Although the phylogenetic distribution is thought to be well characterized (Fig. 1), the origin and evolution of many aspects of DUI remains poorly understood. For example, there are conflicting hypotheses regarding whether male (M) mtDNA has originated once and has been lost multiple times (Stewart *et al*. 2009, 2021; Doucet-Beaupré *et al*. 2010), or if it has originated independently multiple times (Hoeh *et al*. 1996; Maeda *et al*. 2021). Uncertainty stems from inconsistent phylogenetic relationships between female (F) and M mtDNA, and non-monophyly of M mtDNA. Phylogenetic relationships between F and M mtDNA in DUI taxa exhibit two distinct patterns. Female and M mtDNA are reciprocally monophyletic across species in some orders, while they show sister relationships within a species in others. In other words, M mtDNA is non-monophyletic across all DUI species but shows topologies consistent with a single origination in some lineages (Unionida), independent originations in others (Mytilida, Nuculanida, Venerida), or has not been examined in more than one species (Cardiida) in yet others (Breton, Stewart and Blier 2009; Gusman *et al*. 2016). Depending on the lineage, F and M mtDNA genes can be up to 90% identical (Mytilida and Venerida) or differ by more than 50% in their amino acid sequences (Unionida) (Mizi, Zouros and Rodakis 2006; Breton *et al*. 2007; Breton, Stewart and Blier 2009; Gusman *et al*. 2016).

**Figure 1.**
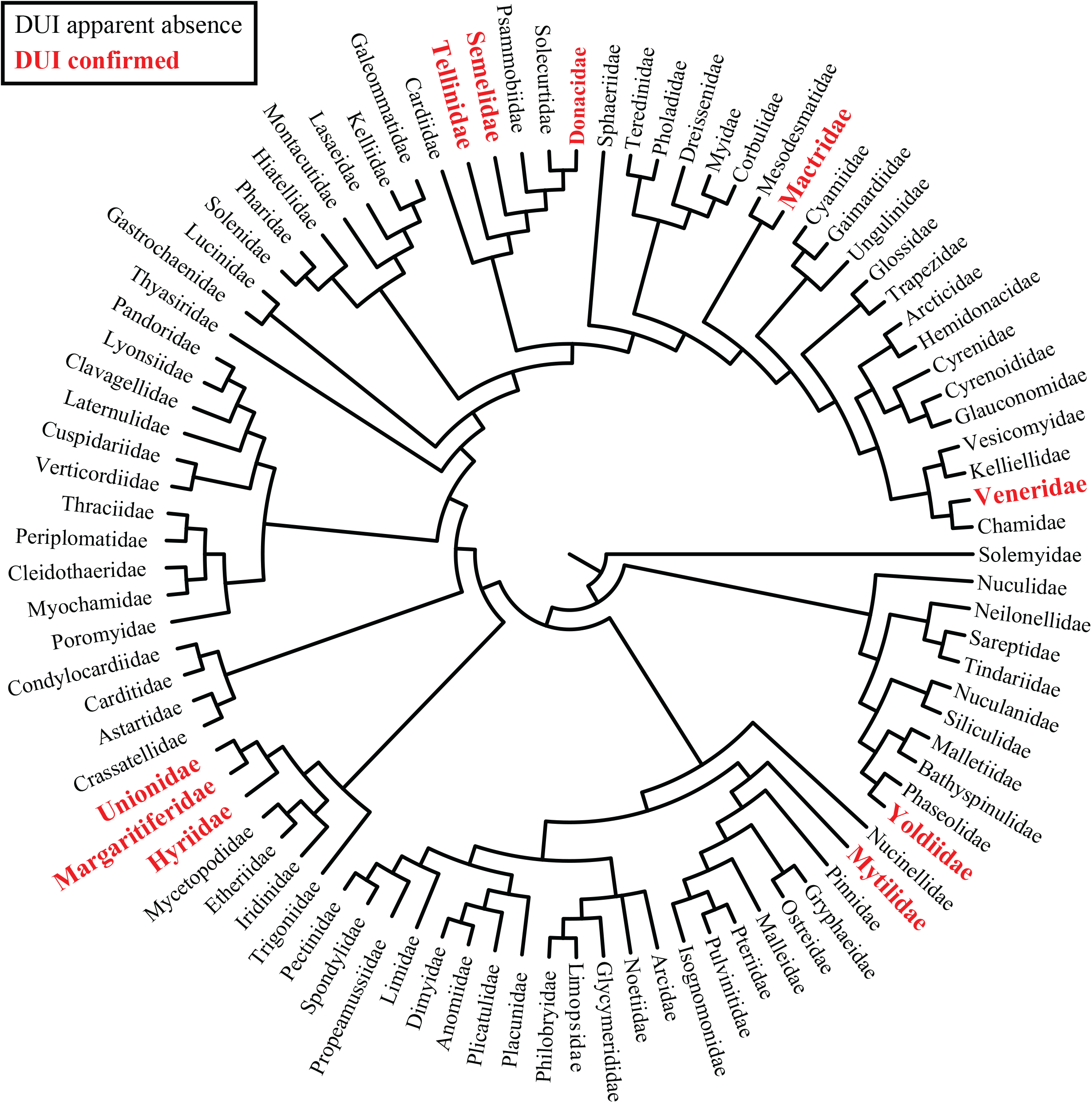
Phylogenetic distribution of doubly uniparental inheritance (DUI) based on a family-level tree of Bivalvia presented in Combosch *et al*. (2017). Families confirmed to exhibit DUI are bolded and colored red.

Recombination events between F and M mtDNA have been documented in several DUI species *(Mytilus* spp. and *Ruditapes philippinarum*) (Ladoukakis and Zouros 2001; Burzyński *et al*. 2003; Passamonti, Boore and Scali 2003; Filipowicz *et al*. 2008; Ladoukakis *et al*. 2011). These events are similar to homologous recombination in bacteria, where novel fragments from the donor genome replace existing homologous genetic material in the recipient genome (Spratt *et al*. 1992). In *Mytilus*, mitochondrial recombination often precipitates a “role-reversal” in which the F mtDNA receives a M control region and is subsequently transmitted as M mtDNA (Cao *et al*. 2004; Mizi, Zouros and Rodakis 2006; Stewart *et al*. 2009; Kyriakou *et al*. 2015). In this event, recombination erases divergence between the rest of the F and M mtDNA genes (e.g., those involved with oxidative phosphorylation (OXPHOS)). This recombination also results in a phylogenetic pattern of concerted evolution in OXPHOS genes, which could cause the observed conflict in sequence divergence and topologies of F and M mtDNA between DUI lineages (Stewart *et al*. 2009; Gusman *et al*. 2016). Recombination events have also been documented to occur in other areas of mtDNA in DUI species (Burzyński *et al*. 2003; Passamonti, Boore and Scali 2003), including within OXPHOS genes (Ladoukakis and Zouros 2001; Ladoukakis *et al*. 2011). If occasional recombination in OXPHOS genes has occurred throughout the evolutionary history of bivalves, certain OXPHOS genes could retain sites informative about the origin of M mtDNA, but signal from these sites has likely been masked when using concatenation-based methods. Recent advances in site-based methodologies that estimate concordance at the level of individual sites, including the site concordance factor (Minh, Hahn and Lanfear 2020), are therefore useful for investigating the origin of M mtDNA.

Mitochondrial recombination is well-documented in Mytilida and Venerida, but recombination is apparently absent in Unionida. This may be due a large extension in the *COX2* gene in the M mtDNA or the presence of sex-specific open reading frames (*orfs*) in the F and M mtDNA (Stewart *et al*. 2009; Breton *et al*. 2011; Gusman *et al*. 2016). Most DUI bivalves exhibit extensions to the *COX2* gene in the M mtDNA, ranging from ~300 bp to 4.5 kb (Curole and Kocher 2002; Bettinazzi, Plazzi and Passamonti 2016; Capt *et al*. 2020), which have been hypothesized to serve as a tag for cells or organelles harboring M mtDNA (Chakrabarti *et al*. 2007). Sex-specific *orfs* likely originated via duplication and have been confirmed to code for proteins in Mytilida, Unionida, and Venerida (Breton *et al*. 2011; Milani *et al*. 2014; Ouimet *et al*. 2020). Although their function is uncertain, it is hypothesized *orfs* are involved in sex determination or sexual development (Breton *et al*. 2011, 2022; Milani *et al*. 2014; Guerra *et al*. 2019; Ouimet *et al*. 2020). Although *COX2* extensions and sex-specific *orfs* are found in most DUI lineages, they have been comparably evolutionarily conserved across Unionida (Curole and Kocher 2002; Guerra *et al*. 2019), suggesting one of these two characteristics may explain why recombination is selected against.

In this study, we revisit the related issues of the origins of M mtDNA and recombination in mtDNA. Specifically, we use phylogenetic methods to 1) investigate the number of origins of M mtDNA, 2) infer the prevalence of mitochondrial recombination, and 3) investigate the potential drivers or inhibitors of mtDNA recombination. Our findings support a single origination of M mtDNA in bivalves with occasional recombination events causing observed non-monophyly of M mtDNA using concatenation-based methods.

## Materials and Methods

### Phylogenetic distribution of doubly uniparental inheritance

To provide an overview of the phylogenetic distribution of DUI in bivalves, we downloaded the phylogeny presented in Combosch *et al*. (2017). We collapsed the phylogeny to the family-level (93 families; see Table S1) and compiled DUI reports from the literature (Theologidis *et al*. 2008; Gusman *et al*. 2016; Capt *et al*. 2020).

### Mitogenomic dataset and phylogenetic analyses

We downloaded M and F mitogenomes for 37 DUI species and 10 representative orders in Bivalvia from the NCBI nucleotide collection (Table S2). *Octopus bimaculatus* (Cephalopoda) was used as an outgroup. In cases where annotations of mitogenomes were incomplete, we used MITOS2 (Bernt *et al*. 2013) to identify protein-coding genes. We excluded *ATP8* due to missing data across most species and a partial portion of *COX2* for *Limecola balthica* and *Scrobicularia plana* (Cardiida) M mtDNA due to a large insertion (Capt *et al*. 2020). Protein-coding genes were aligned using MACSE v 2.05 (Ranwez and Douzery 2018). We then concatenated the 12 mitochondrial genes and removed all sites with missing data. The resulting concatenated alignment was used for phylogenetic analysis and consisted of 83 sequences represented by 2,622 amino acids (File S1). A phylogeny was estimated in IQ-TREE v 2.2.0.3 (Minh *et al*. 2020) using 10 independent runs. ModelFinder (Kalyaanamoorthy *et al*. 2017) was used to select the best amino acid model of evolution (mtInv+F+I+G4) and 10^3^ ultrafast bootstrap replicates were used to assess nodal support (Hoang *et al*. 2018).

We used site concordance factors (Minh, Hahn and Lanfear 2020) to test M mtDNA origination hypotheses. Briefly, site concordance factors measure the percentage of sites supporting a certain branch in a phylogeny. Hypotheses can be tested by comparing observed site concordance factors with a distribution of site concordance factors from data simulated under a given phylogenetic hypothesis (e.g., Hibbins, Gibson and Hahn 2020). We used site concordance factors from both individual genes and a concatenated alignment of all genes to test two hypotheses: 1) ten independent originations of M mtDNA (as supported by concatenation methods; Fig. 2), and 2) a single origination of M mtDNA. Specifically, our methodology evaluated these two hypotheses by directly comparing observed site concordance factors for a single origination of M mtDNA to a distribution of site concordance factors for a single origination of M mtDNA that could occur by chance under multiple origins. To generate distributions of site concordance factors for hypothesis testing from the concatenated dataset and each gene independently, we used AliSim (Ly-Trong *et al*. 2022) to simulate 10^3^ amino acid datasets based on the resolved topology from each empirical alignment using the best model of amino acid evolution as determined by ModelFinder. We chose to use AliSim over other methods (e.g., Seq-Gen, Dawg, INDELible) to account for the non-independence of mtDNA substitutions. Next, we used Mesquite v 3.3.1 (Maddison and Maddison 2017) to create a topology from the concatenated analysis that enforced the monophyly of all M mtDNA while retaining branch length information (Fig. S1; File S2). We then calculated site concordance factors for all empirical and simulated datasets using 100 quartets. With those, we gathered site concordance factors for the branch coinciding to a single origin of M mtDNA (Fig. S1) and used one-tailed tests (with p = 0.05) to determine if the observed site concordance factor was significantly larger than expected under 10 independent originations.

**Figure 2.**
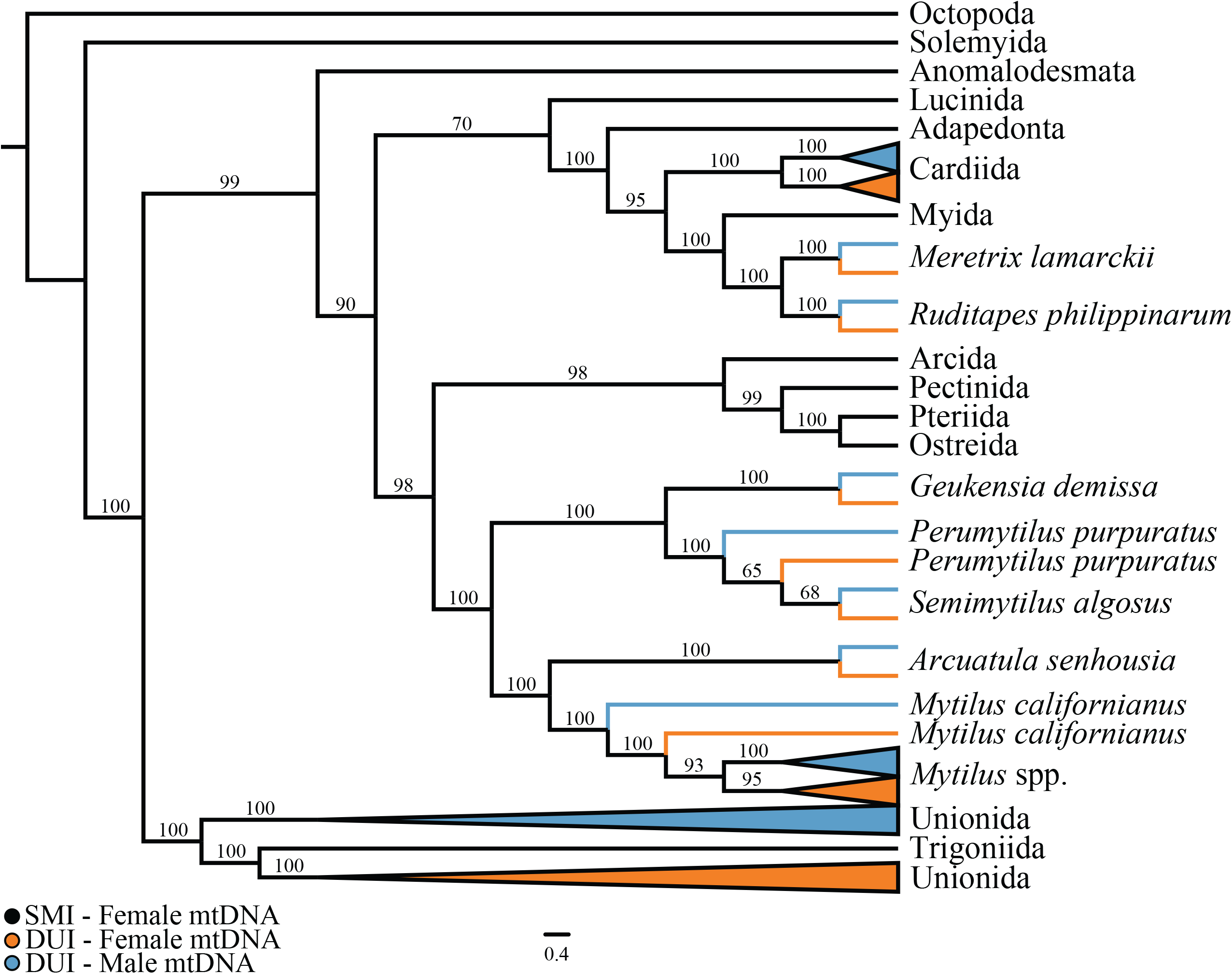
Phylogeny of the class Bivalvia based on amino acid sequences for 12 mitochondrial genes, showing lineages with strictly maternal inheritance (SMI), female mtDNA in DUI species, and male mtDNA in DUI species. *Mytilus* spp. refers to *M. edulis, M. galloprovincialis*, and *M. trossolus*. Values above branches represent ultrafast bootstrap support.

We investigated the hypothesis that the lack of recent recombination in Cardiida and Unionida is a result of intensified selection on M mtDNA genes that have adapted to male functions. We chose to perform this test in Cardiida given we resolved a similar phylogenetic pattern between F and M mtDNA as Unionida (Fig. 2). We used RELAX (Wertheim *et al*. 2015) in HyPhy v 2.5.25 (Pond, Frost and Muse 2005) with a concatenated nucleotide alignment of 12 M mtDNA genes (File S3; Table S3) to test if selection on M mtDNA in Cardiida and Unionida was significantly different than Mytilida and Venerida. Considering extensions to the *COX2* gene in the M mtDNA are shared in Cardiida and Unionida and hypothesized to be a proximate cause of the absence of recombination, we also used RELAX independently on a nucleotide alignment of M mtDNA *COX2* gene (File S4; Table S3). Codons with missing or ambiguous data in each alignment were removed. Likelihood ratio tests were used to evaluate models with a significance level of p = 0.05.

### Estimation of recombination frequency

To estimate the frequency of recombination, we estimated divergence times between F and M mtDNA lineages. We used BEAST v 2.6.7 (Bouckaert *et al*. 2019) with a concatenated nucleotide alignment of 12 F and M mtDNA genes for all taxa sampled in Mytilida (File S5; Table S4), where recombination between M and F mtDNA has been observed and reliable fossil calibrations are available. Codons with missing or ambiguous data in each alignment were removed. The best fit model of nucleotide evolution for each codon position was selected by ModelFinder, a relaxed molecular clock was fit to each codon position, and a calibrated Yule process was used as the tree prior. We enforced priors that date the MRCA of F and M mtDNA for *Mytilus edulis, M. galloprovincialis*, and *M. trossolus* between 3.1 and 4.8 Mya (Rawson and Harper 2009). The analysis was run for 10^8^ MCMC generations with an initial 10% burn-in. Tracer v1.7.1 (Rambaut *et al*. 2018) was used to determine the appropriate burn-in value and ensure convergence of all parameters (ESS > 200), and a maximum clade credibility tree was created using TREEANNOTATOR v 2.6 (Bouckaert *et al*. 2019). To get a rough estimate of the timing of recombination events, we calculated an average divergence time between putatively recombinant F and M mtDNA lineages.

## Results and Discussion

Phylogenetic reconstruction based on the concatenated alignment of 12 of the 13 mitochondrial protein coding OXPHOS genes showed non-monophyly of M mtDNA across bivalves (Fig. 2; File S6), as shown previously (Hoeh *et al*. 1996; Gusman *et al*. 2016; Maeda *et al*. 2021). While this topology has been previously interpreted as consistent with multiple origins or losses of M mtDNA (Hoeh *et al*. 1996; Stewart *et al*. 2009; Doucet-Beaupré *et al*. 2010; Gusman *et al*. 2016; Maeda *et al*. 2021), it is also consistent with concerted evolution due to recombination between F and M mtDNA in Mytilida and Venerida, and a lack of recombination in Cardiida and Unionida (Fig. 2). As has been seen in previous studies (Gusman *et al*. 2016; Maeda *et al*. 2021), we found that F and M mtDNA within species in Mytilida and Venerida are generally sister, which is expected under the hypothesis of recombination between F and M mtDNA. One exception is *Mytilus edulis, M. galloprovincialis*, and *M. trossolus (Mytilus* spp.) have reciprocally monophyletic F and M mtDNA (Fig. 2), despite the fact that *Mytilus* spp. are known to recombine (Ladoukakis and Zouros 2001; Burzyński *et al*. 2003; Filipowicz *et al*. 2008; Ladoukakis *et al*. 2011). We estimate that recombinant M mtDNA fix less frequently (~11 My; 95% CI: 7.3-14.5 My; Fig. S2) than do speciation events in *Mytilus* (~3.1-4.8 Mya). We hypothesize the reciprocal monophyly of F and M mtDNA will appear frequently across the phylogeny of certain DUI bivalve lineages at shallow taxonomic scales when data for additional taxa become available.

Mitochondrial recombination in Mytilida and Venerida results in a pattern of concerted evolution of F and M mtDNA, which may be favored to combat the deleterious effects of asexual inheritance and maintain mitonuclear compatibility across tissues (Muller 1964). If there are two sets of highly divergent mtDNAs within the same organism, interacting nuclear genes necessary for proper function may not cooperate efficiently with both mtDNAs, resulting in mitonuclear incompatibility for one mitogenome (Hill 2015). Mitonuclear coevolution has recently been confirmed in bivalves, with highly correlated evolution between mitochondrial and nuclear subunits involved with OXPHOS (Piccinini *et al*. 2021). However, relaxed selection on M mtDNA may be common in DUI bivalves, therefore favoring nuclear coevolution with F over M mtDNAs (Maeda *et al*. 2021). Here we suggest that mitonuclear compatibility may be restored via recombination in some DUI lineages in an analogous process to the “Fountain of Youth” (Perrin 2009). In this process, occasional recombination events are hypothesized to counteract accumulated deleterious mutations in previously non-recombining sex chromosomes (Perrin 2009).

Analyses of energetic metabolism provide support that mitochondrial recombination may be favored to purge deleterious mutations in M mtDNA. In Mytilida and Venerida, sperm are dependent on OXPHOS to sustain motility (Bettinazzi *et al*. 2019, 2020), which highlights the importance of compatibility between M mtDNA and nuclear genes. Comparative physiological studies in *M. edulis* have shown that recombination events do not have obvious deleterious effects on sperm performance (Everett *et al*. 2004). Rather, recombination may be advantageous because sperm with recently masculinized M mtDNA (i.e., those carrying F mtDNA with M control regions) swim faster than those with ancestral M mtDNA (Jha *et al*. 2007). Sperm swimming velocity has been demonstrated to be correlated with ATP levels in many taxa (Perchec *et al*. 1995; Burness, Moyes and Montgomerie 2005), and ATP production is lower in sperm with M mtDNA than eggs with F mtDNA (Bettinazzi *et al*. 2019). Mitochondrial recombination, therefore, may be favored to maximize M mtDNA ATP production in Mytilida and Venerida by replacing defective M mtDNA OXPHOS genes with more energetically robust F mtDNA OXPHOS genes (Breton, Stewart and Blier 2009). To our knowledge, physiological studies have been limited to Mytilida and Venerida (Bettinazzi *et al*. 2020), and future analogous studies in Cardiida and Unionida may further support our hypothesis.

We find a different pattern of phylogenetic relationships of mtDNAs in Unionida when compared to Mytilida and Venerida, consistent with previous studies (Gusman *et al*. 2016). In Unionida, F and M mtDNA are reciprocally monophyletic across species (Fig. 2). A similar relationship was recovered in Cardiida (Fig. 2), albeit based on two species. However, *L. balthica* (Cardiida: Tellinidae) and *S. plana* (Cardiida: Semelidae) are estimated to have diverged at or near the Cretraceous-Palogene boundary (~66 Mya) (Crouch *et al*. 2021), far greater than our estimated frequency of recombinant fixation in Mytilida (~11 My). Therefore, our data is consistent with the absence of recent recombination between F and M mtDNA in both Cardiida and Unionida. We hypothesize mitochondrial recombination was the plesiomorphic condition of DUI species and was independently lost in these lineages. This is because M mtDNA in Cardiida and Unionida would be monophyletic had recombination independently originated in Mytilida and Venerida. One possible explanation for the loss of recombination in Cardiida and Unionida involves a large extension of *COX2* in the M mtDNA (Curole and Kocher 2002), which is hypothesized to promote gender-specific mitochondrial localizations (Chakrabarti *et al*. 2007). Recombination between F and M mtDNA could disrupt proper localization and therefore be selected against.

Although large extensions to *COX2* may be a proximate cause for the loss of recombination in Cardiida and Unionida, its adaptive significance remains unclear. If *COX2* or additional M mtDNA genes are adapted to certain male functions, those adapted features could be lost following recombination with F mtDNA. Were this the case, we might expect to see intensified selection on *COX2* and M mtDNA genes in Cardiida and Unionida compared to Mytilida and Venerida. Our analyses reject this hypothesis, and in fact indicate significant evidence of relaxed selection in Cardiida and Unionida (*COX2*: K = 0.71, p = 0.001; 12 genes: K = 0.44, p < 0.001; Table S5). Another possible explanation for the loss of recombination is that mtDNA may have a role in sex determination, particularly in Unionida (Breton *et al*. 2011). Unlike other bivalve lineages with DUI, some families in Unionida (i.e., Margaritiferidae and Unionidae) have evolutionarily conserved sex-specific *orfs* (F-*orf* and M-*orf*) that have been confirmed to code for proteins (Breton *et al*. 2011). Additionally, hermaphroditism has evolved multiple times in these lineages, and each transition is often associated with the origin of a F-like mtDNA that has a hermaphrodite-specific *orf* (Breton *et al*. 2011 but see Soroka and Burzyński 2017). This suggests mtDNA *orfs* are associated with sexual transitions in Unionida and may have a role in sex determination or sexual development (Breton *et al*. 2011, 2014, 2022). Recombination between F and M mtDNA would therefore be deleterious, albeit we recognize this explanation may be limited to the families Margaritiferidae and Unionidae.

In principle, gene trees could be used to determine the number of origins of M mtDNA. In the absence of mitochondrial recombination, a single origin of M mtDNA would result in reciprocal monophyly of F and M mtDNA across DUI species. However, it is unlikely that gene trees with the appropriate topology will be observed when there is recombination. Therefore, our phylogenetic reconstruction (Fig. 2) is consistent with either multiple origins of M mtDNA (up to 10) or a single origination of M mtDNA with recombination acting in a lineage-specific manner over long evolutionary timescales. We tested these hypotheses using site concordance factors, which supported a single origination of M mtDNA followed by lineage-specific recombination (Fig. 3; Table S6). Specifically, we found more site-level support for a single origin and can reject multiple origin hypotheses using both an individual OXPHOS gene *(ND1:* p=0.03; Fig. 3; Table S6) and a concatenated alignment of 12 genes (p < 0.001; Fig. 3; Table S6). Our results agree with hypotheses presented in previous studies (Hoeh *et al*. 1997; Theologidis *et al*. 2008; Stewart *et al*. 2009; Doucet-Beaupré *et al*. 2010; Zouros 2013).

**Figure 3.**
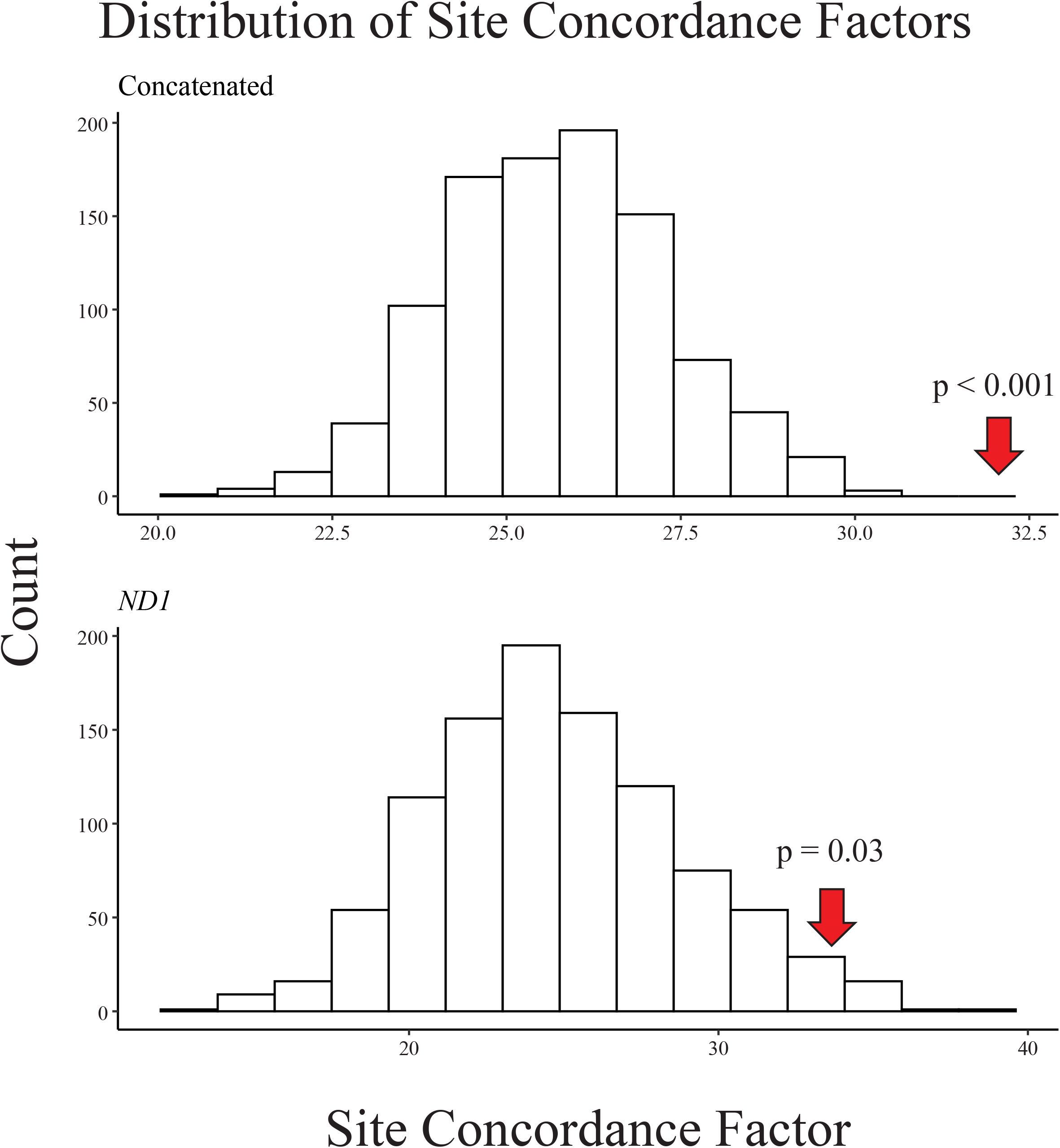
Null distribution and observed site concordance factors used to assess support for a single origination of male mitochondrial DNA for a concatenated alignment and *ND1*. In each plot, white bars represent the null distribution based on 1000 simulated amino acid datasets, the red arrow represents the observed value based on empirical data, and the p-value is reported.

Although we can reject multiple origination hypotheses, the retention of signal in protein coding genes for a single origin of M mtDNA remains unclear. Recombination events have been documented to occur throughout mtDNA in DUI species, including within mitochondrial genes, but have been hypothesized to only occur in somatic tissue and not inherited through gametes (Ladoukakis and Zouros 2001; Ladoukakis *et al*. 2011). Given this context, our results suggest this conclusion may be unrealistic. Future investigations across DUI bivalves may reveal more complex patterns of recombination in protein coding genes and inheritance of recombinant mtDNAs, which could explain preserved signal for a single origination of M mtDNA in mitochondrial OXPHOS genes.

## Conclusion

Our results support a single origination of M mtDNA followed by lineage-specific recombination, which has led to non-monophyly of M mtDNA using concatenation-based methods. Mitochondrial recombination events may occur to counteract the accumulation of deleterious mutations in M mtDNA to restore ATP production but are exclusive to Mytilida and Venerida (based on available data). It remains uncertain why recombination is absent in Cardiida and Unionida, but it may be selected against because of the role of mtDNAs in sex determination or sexual development in these lineages. Future studies into these topics will further contribute to the understanding of DUI and the functional significance of retaining M mtDNA in bivalves.

## Supporting information

All supplemental figures, files, and tables

## Data Availability Statement

Data used in this study can be found on GenBank with all accession numbers used as part of this research found in Supplementary Materials.

## Acknowledgments

The authors wish to thank the Havird and Kirkpatrick lab groups for discussions regarding this research. This work was supported by funds awarded to Chase Smith from the University of Texas at Austin Stengl-Wyer Endowment, NIH grant R35-GM142836 awarded to Justin Havird, and NIH grant R01-GM116853 awarded to Mark Kirkpatrick.

## Notes

### Competing Interest Statement

The authors have declared no competing interest.

