## Supplementary material for "A tale of two paths: The evolution of mitochondrial recombination in bivalves with doubly uniparental inheritance": All supplemental figures, files, and tables: Supplemental Materials.pdf

**Supplemental Tables**

**Table S1.** Families and tip labels from Combasch et al. (2017) used to evaluate the phylogenetic distribution of doubly uniparental inheritance.

| Family | Tip Label |
|---|---|
| Anomiidae | Anomia_ephippium |
| Arcidae | Arca_noae |
| Arcticidae | Arctica_islandica |
| Astartidae | Astarte_sulcata |
| Bathyspinulidae | Bathyspinula_hilleri |
| Cardiidae | Tridacna_maxima |
| Carditidae | Cardita_calyculata |
| Chamidae | Chama_asperella |
| Clavagellidae | Clavagella_melitensis |
| Cleidothaeridae | Cleidothaerus_albidus |
| Condylocardiidae | Carditella_capensis |
| Corbulidae | Notocorbula_tunicata |
| Crassatellidae | Crassinella_lunulata |
| Cuspidariidae | Cuspidaria_cuspidata |
| Cyamiidae | Cyamiomactra_laminifera |
| Cyrenidae | Corbicula_fluminea |
| Cyrenoididae | Cyrenoida_floridana |
| Dimyidae | Dimya_lima |
| Donacidae | Donax_trunculus |
| Dreissenidae | Dreissena_polymorpha |
| Etheriidae | Etheria_elliptica |
| Gaimardiidae | Gaimardia_trapezina |
| Galeommatidae | Galeomma_turtoni |
| Gastrochaenidae | Gastrochaena_dubia |
| Glauconomidae | Glauconome_rugosa |
| Glossidae | Glossus_humanus |
| Glycymerididae | Glycymeris_glycymeris |
| Gryphaeidae | Hyotissa_hyotis |
| Hemidonacidae | Hemidonax_pictus |
| Hiatellidae | Hiatella_arctica |
| Hyriidae | Hyridella_australis |
| Iridinidae | Aspatharia_pfeifferiana |
| Isognomonidae | Isognomon_alatus |
| Kelliellidae | Kelliella_cfnitida |
| Kelliidae | Bornia_sebetia |
| Lasaeidae | Lasaea_adansoni |

| | |
|---|---|
| Laternulidae | Laternula_elliptica |
| Limidae | Lima_lima |
| Limopsidae | Limopsis_sp |
| Lucinidae | Lucina_pensylvanica |
| Lyonsiidae | Lyonsia_floridana |
| Mactridae | Mactra_violacea |
| Malleidae | Malleus_albus |
| Malletiidae | Malletia_johnsoni |
| Margaritiferidae | Margaritifera_margaritifera |
| Mesodesmatidae | Donacilla_cornea |
| Montacutidae | Mysella_charcoti |
| Mycetopodidae | Anodontites_elongata |
| Myidae | Mya_arenaria |
| Myochamidae | Myadora_brevis |
| Mytilidae | Mytilus_edulis |
| Neilonellidae | Neilonella_whoii |
| Noetiidae | Eontia_ponderosa |
| Nucinellidae | Nucinella_giribeti |
| Nuculanidae | Nuculana_pernula |
| Nuculidae | Nucula_atacellana |
| Ostreidae | Ostrea_edulis |
| Pandoridae | Pandora_pinna |
| Pectinidae | Pecten_maximus |
| Periplomatidae | Cochlodesma_praetenue |
| Pharidae | Phaxas_pellucidus |
| Phaseolidae | Lametila_abyssorum |
| Philobryidae | Philobrya_sublaevis |
| Pholadidae | Pholas_dactylus |
| Pinnidae | Pinna_carnea |
| Placunidae | Placuna_placenta |
| Plicatulidae | Plicatula_plicata |
| Poromyidae | Poromya_illevis |
| Propeamussiidae | Propeamussium_sp |
| Psammobiidae | Gari_maculosa |
| Pteriidae | Pteria_hirundo |
| Pulvinitidae | Pulvinites_exempla |
| Sareptidae | Pristigloma_nitens |
| Semelidae | Abra_alba |
| Siliculidae | Silicula_sp |
| Solecurtidae | Tagelus_plebeius |
| Solemyidae | Solemya_velesiana |
| Solenidae | Solen_vaginoides |
| Sphaeriidae | Sphaerium_nucleus |

| | |
|---|---|
| Spondylidae | Spondylus_ambiguus |
| Tellinidae | Angulus_versicolor |
| Teredinidae | Teredo_clappi |
| Thraciidae | Thracia_villosiuscula |
| Thyasiridae | Thyasira_equalis |
| Tindariidae | Tindaria_kennerlyi |
| Trapezidae | Trapezium_sublaevigatum |
| Trigoniidae | Neotrigonia_margaritacea |
| Ungulinidae | Cycladicama_cumingi |
| Unionidae | Unio_pictorum |
| Veneridae | Venus_verrucosa |
| Verticordiidae | Haliris_fischeriana |
| Vesicomyidae | Calyptogena_magnifica |
| Yoldiidae | Yoldia_limatula |

**Table S2.** GenBank accessions for female (F) and male (M) mitochondrial genomes used in phylogenetic analyses of 12 OXPHOS genes.

| **Order** | **Taxon** | **Accession F** | **Accession M** |
|---|---|---|---|
| Adapedonta | *Solen grandis* | HQ703012 | - |
| Anomalodesmata | *Myadora brevis* | KX815961 | - |
| Arcida | *Arca navicularis* | MG641752 | - |
| Cardiida | *Limecola balthica* | MN528028 | MN528029 |
| | *Scrobicularia plana* | MN528026 | MN528027 |
| Lucinida | *Lucinella divaricata* | EF043342 | - |
| Myida | *Dreissena rostriformis* | MW080914 | - |
| Mytilida | *Arcualata senhousia* | GU001953 | GU001954 |
| | *Geukensia demissa* | MN449487 | MN449488 |
| | *Mytilus californianus* | JX486124 | JX486123 |
| | *Mytilus edulis* | MF407676 | AY823623 |
| | *Mytilus galloprovincialis* | FJ890849 | FJ890850 |
| | *Mytilus trossulus* | GU936625 | GQ438250 |
| | *Perumytilus purpuratus* | MH330333 | MH330330 |
| | *Semimytilus algosus* | MT026712 | MT026713 |
| Octopoda | *Octopus bimaculatus* | KT581981 | - |
| Ostreida | *Crassostrea gigas* | EU672831 | - |
| Pectinida | *Pecten maximus* | KP900975 | - |
| Pteriida | *Pinctada margaritifera* | HM467838 | - |
| Solemyida | *Petrasma pervernicosa* | KY244080 | - |

| | | | |
|---|---|---|---|
| Trigoniida | *Neotrigonia margaritacea* | KU873118 | - |
| Unionida | *Anodonta anatina* | KF030964 | KF030963 |
| | *Arconaia lanceolata* | KJ144818 | KJ775864 |
| | *Chamberlainia hainesiana* | MK994770 | MK994771 |
| | *Cumberlandia monodonta* | KU873123 | KU873124 |
| | *Echyridella menziesii* | KU873121 | KU873122 |
| | *Lampsilis siliquoidea* | MF326973 | MF326974 |
| | *Lampsilis powelli* | MF326971 | MF326972 |
| | *Margaritifera margaritifera* | MK421956 | MK421959 |
| | *Microcondylaea bonellii* | MK994772 | MK994773 |
| | *Monodontina vondembuschiana* | MK994774 | MK994775 |
| | *Pilsbryoconcha exilis* | MK994776 | MK994777 |
| | *Potamida littoralis* | KT247374 | KT247375 |
| | *Potamilus alatus* | KU559011 | KU559010 |
| | *Pyganodon grandis* | FJ809754 | FJ809755 |
| | *Quadrula quadrula* | FJ809750 | FJ809751 |
| | *Sinanodonta woodiana* | HQ283346 | MH349356 |
| | *Solenaia carinata* | KC848654 | KC848655 |
| | *Solenaia oleivora* | KF296320 | KY007143 |
| | *Unio crassus* | KY290447 | KY290450 |
| | *Unio delphinus* | KT326917 | KT326918 |
| | *Unio pictorum* | HM014134 | MH349358 |
| | *Unio tumidus* | KY021076 | KY021075 |
| | *Utterbackia peninsularis* | HM856636 | HM856635 |
| | *Venustaconcha ellipsiformis* | FJ809753 | FJ809752 |
| Venerida | *Meretrix larmarckii* | KP244451 | KP244452 |
| | *Ruditapes philippinarum* | AB065375 | AB065374 |

**Table S3.** GenBank accession numbers for male mitochondrial genomes used in RELAX analyses.

| **Order** | **Taxon** | **Accession** |
|---|---|---|
| Cardiida | *Limecola balthica* | MN528029 |
| | *Scrobicularia plana* | MN528027 |
| Mytilida | *Arcualata senhousia* | GU001954 |
| | *Geukensia demissa* | MN449488 |
| | *Mytilus californianus* | JX486123 |

| | | |
|---|---|---|
| | *Mytilus edulis* | AY823623 |
| | *Mytilus galloprovincialis* | FJ890850 |
| | *Mytilus trossulus* | GQ438250 |
| | *Perumytilus purpuratus* | MH330330 |
| | *Semimytilus algosus* | MT026713 |
| Unionida | *Anodonta anatina* | KF030963 |
| | *Arconaia lanceolata* | KJ775864 |
| | *Chamberlainia hainesiana* | MK994771 |
| | *Cumberlandia monodonta* | KU873124 |
| | *Echyridella menziesii* | KU873122 |
| | *Lampsilis powelli* | MF326972 |
| | *Lampsilis siliquoidea* | MF326974 |
| | *Margaritifera margaritifera* | MK421959 |
| | *Microcondylaea bonellii* | MK994773 |
| | *Monodontina vondembuschiana* | MK994775 |
| | *Pilsbryoconcha exilis* | MK994777 |
| | *Potamida littoralis* | KT247375 |
| | *Potamilus alatus* | KU559010 |
| | *Pyganodon grandis* | FJ809755 |
| | *Quadrula quadrula* | FJ809751 |
| | *Sinanodonta woodiana* | MH349356 |
| | *Solenaia carinata* | KC848655 |
| | *Solenaia oleivora* | KY007143 |
| | *Unio crassus* | KY290450 |
| | *Unio delphinus* | KT326918 |
| | *Unio pictorum* | MH349358 |
| | *Unio tumidus* | KY021075 |
| | *Utterbackia peninsularis* | HM856635 |
| | *Venustaconcha ellipsiformis* | FJ809752 |
| Venerida | *Meretrix larmarckii* | KP244452 |
| | *Ruditapes philippinarum* | AB065374 |

**Table S4.** GenBank accession numbers for female (F) and male (M) mitochondrial genomes used in the divergence time analysis of Mytilida.

| Order | Taxon | Accession F | Accession M |
|---|---|---|---|

| | | | |
|---|---|---|---|
| Mytilida | *Arcualata senhousia* | GU001953 | GU001954 |
| | *Geukensia demissa* | MN449487 | MN449488 |
| | *Mytilus californianus* | JX486124 | JX486123 |
| | *Mytilus edulis* | MF407676 | AY823623 |
| | *Mytilus galloprovincialis* | FJ890849 | FJ890850 |
| | *Mytilus trossulus* | GU936625 | GQ438250 |
| | *Perumytilus purpuratus* | MH330333 | MH330330 |
| | *Semimytilus algosus* | MT026712 | MT026713 |

**Table S5.** Omega (ω) values for Cardiida+Unionida and Mytilida+Venerida based on alternative models generated by RELAX from a concatenated alignment of 12 male mitochondrial genes and the male mitochondrial gene *COX2*. Percentages represent the proportion of each alignment assigned to ω values. Both analyses strongly supported ($p \leq 0.0001$) relaxed selection in Cardiida+Unionida.

| **Dataset** | **Group** | $\omega_1$ | $\omega_2$ | $\omega_3$ |
|---|---|---|---|---|
| Concatenated | Cardiida+Unionida | 0.0008 (66.69%) | 0.349 (27.07%) | 2.558 (6.24%) |
| | Mytilida+Venerida | $8.66e^{-8}$ (66.69%) | 0.089 (27.07%) | 8.636 (6.24%) |
| *COX2* | Cardiida+Unionida | 0.015 (82.88%) | 0.618 (16.77%) | 11932049 (0.3%) |
| | Mytilida+Venerida | 0.003 (82.88%) | 0.506 (16.77%) | 9999999171 (0.3%) |

**Table S6.** Empirical average site concordance factor and mean and standard deviation (SD) for simulated average site concordance factors (sCFs) for modeling a single origination of doubly uniparental inheritance. Statistics for 12 genes concatenated and each gene, as well as associated p-values for one-tailed tests are reported. Bolded values were determined to be statically significant.

| **Dataset** | **Empirical** | **Mean** | **SD** | **p** |
|---|---|---|---|---|
| **Concatenated** | **32.06** | **25.72** | **1.59** | **< 0.001** |
| *ATP6* | 27.51 | 25.66 | 4.79 | 0.70 |
| *COX1* | 37.84 | 29.91 | 4.55 | 0.08 |
| *COX2* | 38.18 | 28.23 | 5.53 | 0.07 |
| *COX3* | 43.97 | 40.01 | 5.64 | 0.48 |

| | | | | |
|---|---|---|---|---|
| *CYTB* | 29.21 | 24.97 | 4.09 | 0.30 |
| ***ND1*** | **33.64** | **24.76** | **4.09** | **0.03** |
| *ND2* | 23.91 | 24.20 | 4.37 | 1 |
| *ND3* | 31.36 | 24.50 | 6.09 | 0.26 |
| *ND4* | 42.48 | 23.02 | 3.07 | 0.28 |
| *ND4L* | 36.92 | 31.83 | 8.15 | 0.53 |
| *ND5* | 32.07 | 29.35 | 3.64 | 0.45 |
| *ND6* | 20.79 | 18.49 | 5.22 | 0.66 |

### Supplemental Figures

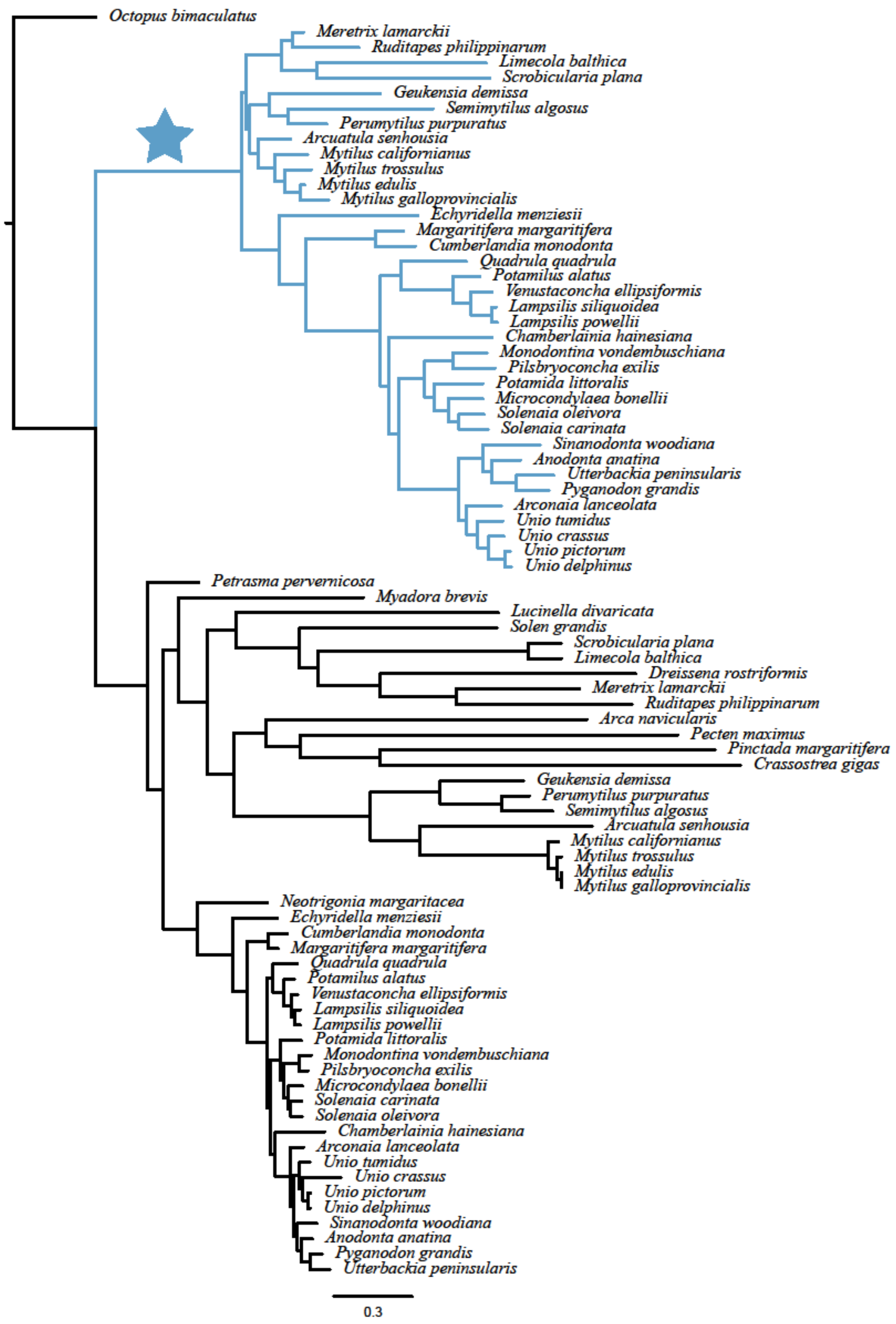


**Figure S1.** Constraint tree enforcing monophyly of male mitochondria used for phylogenetic modeling. Blue branches correspond the male mitochondrial lineages, and the blue star corresponds to the branch for origination testing using site concordance factors.

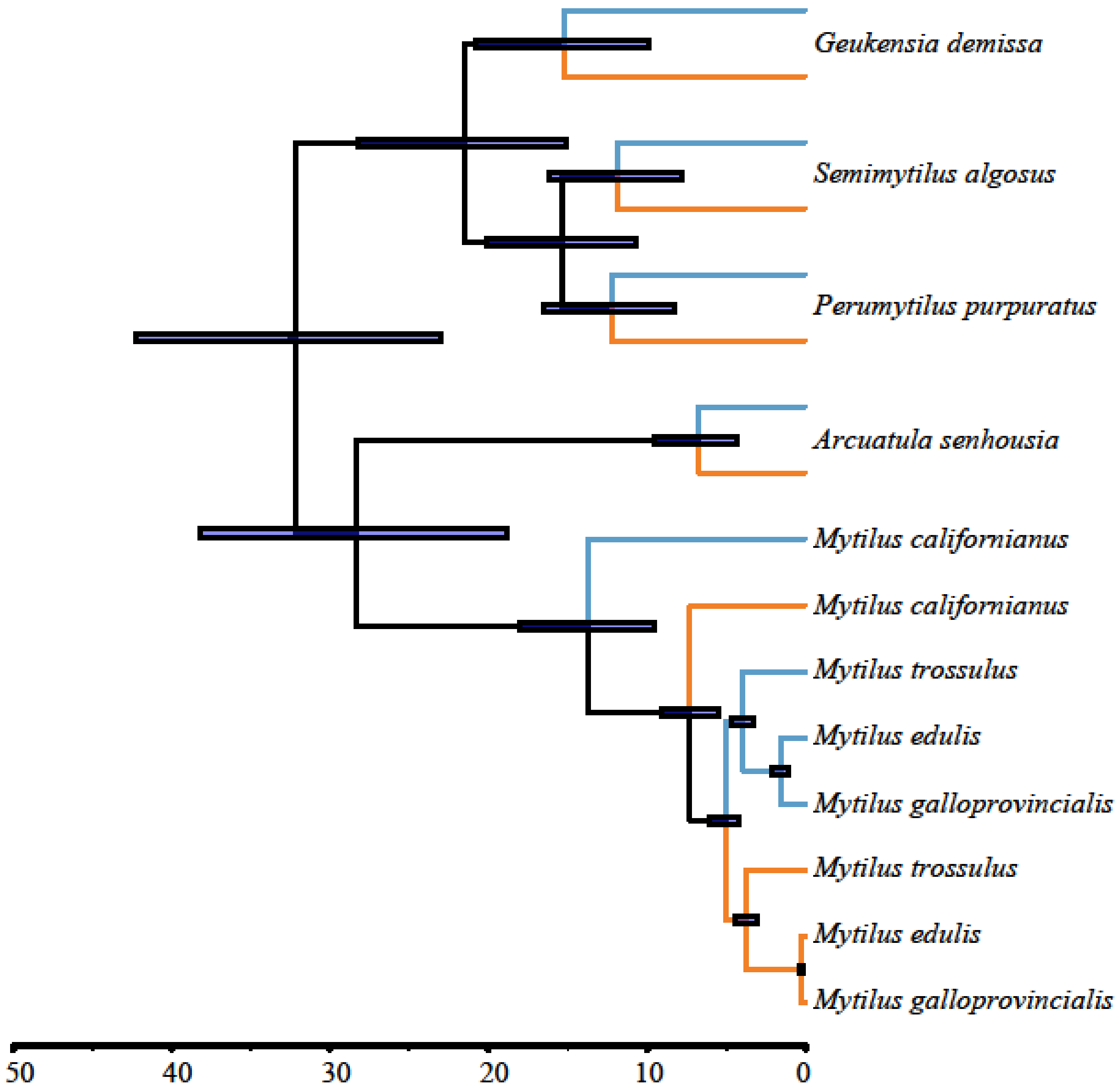


**Figure S2.** Maximum clade credible tree generated from the calibrated phylogenetic analysis of Mytilida in BEAST. Divergence time is scaled to million years before present and node bars represent the 95% CI. Branches representing female and male specific mitochondria are highlighted orange and blue, respectively.

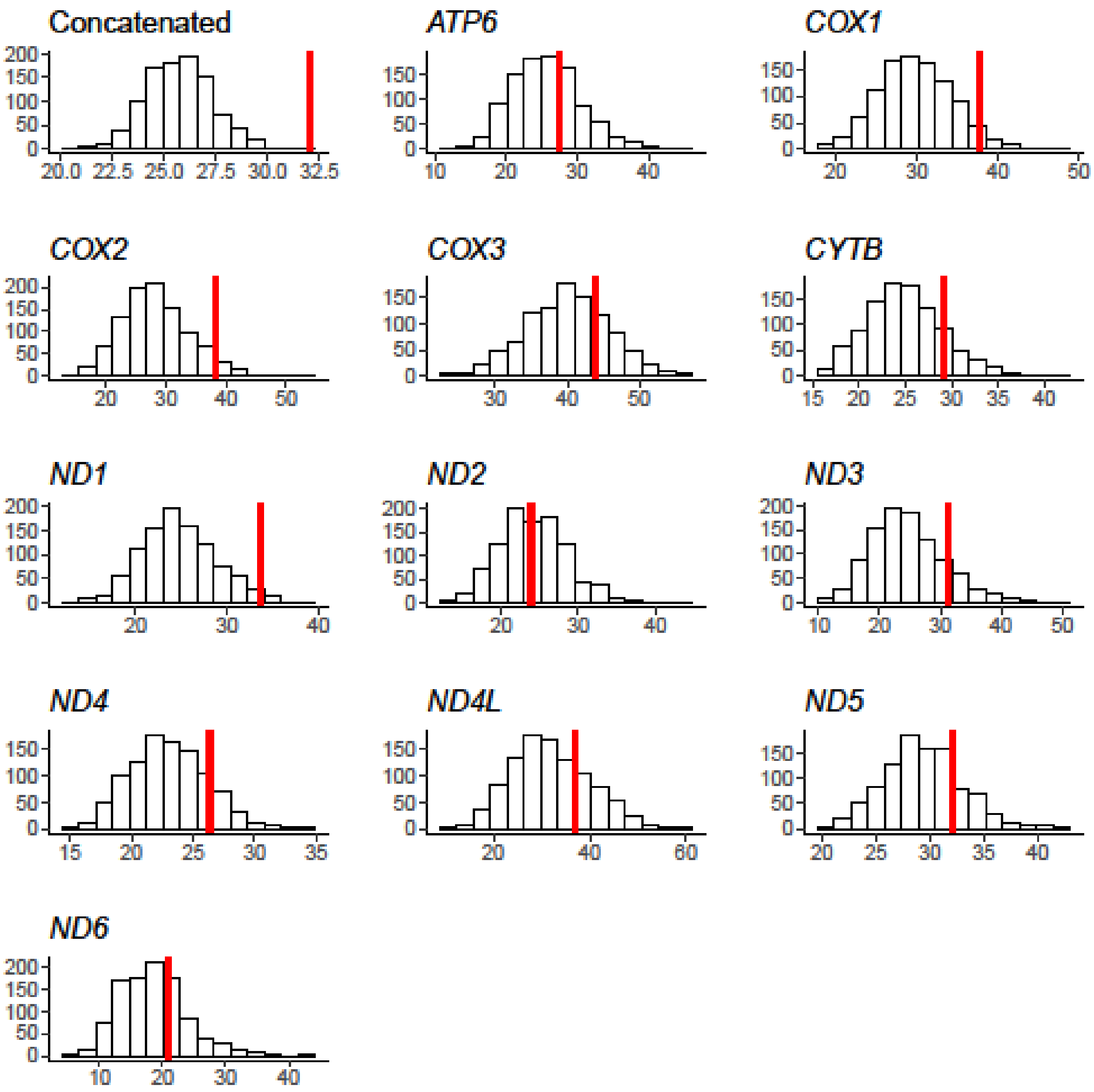


**Figure S3.** Null distribution and observed site concordance factors used to assess support for a single origination of male mitochondrial DNA using a concatenated alignment of 12 gene and each gene independently. In each plot, white bars represent the null distribution based on 1000 simulated amino acid datasets and the red line represents the observed value based on empirical data.

**Supplemental Files**

**File S1.** Concatenated amino acid alignment of 12 mitochondrial genes used in this study in fasta format.

**File S2.** Constraint tree enforcing monophyly of male mitochondria used for phylogenetic modeling.

**File S3.** Concatenated nucleotide alignment of 12 male mitochondrial genes used in the RELAX analysis in fasta format.

**File S4.** Nucleotide alignment of the male mitochondrial gene *COX2* used in the RELAX analysis in fasta format.

**File S5.** Concatenated nucleotide alignment of 12 mitochondrial genes for all Mytilida used in the BEAST analysis in nexus format.

**File S6.** Phylogram generated from a concatenated amino acid alignment of 12 genes in IQ-TREE. Node labels are support values from $10^3$ ultrafast bootstrap replicates.
