## Supplementary figures and images for "A tale of two paths: The evolution of mitochondrial recombination in bivalves with doubly uniparental inheritance"

### FigS1_Constraint_tree.pdf

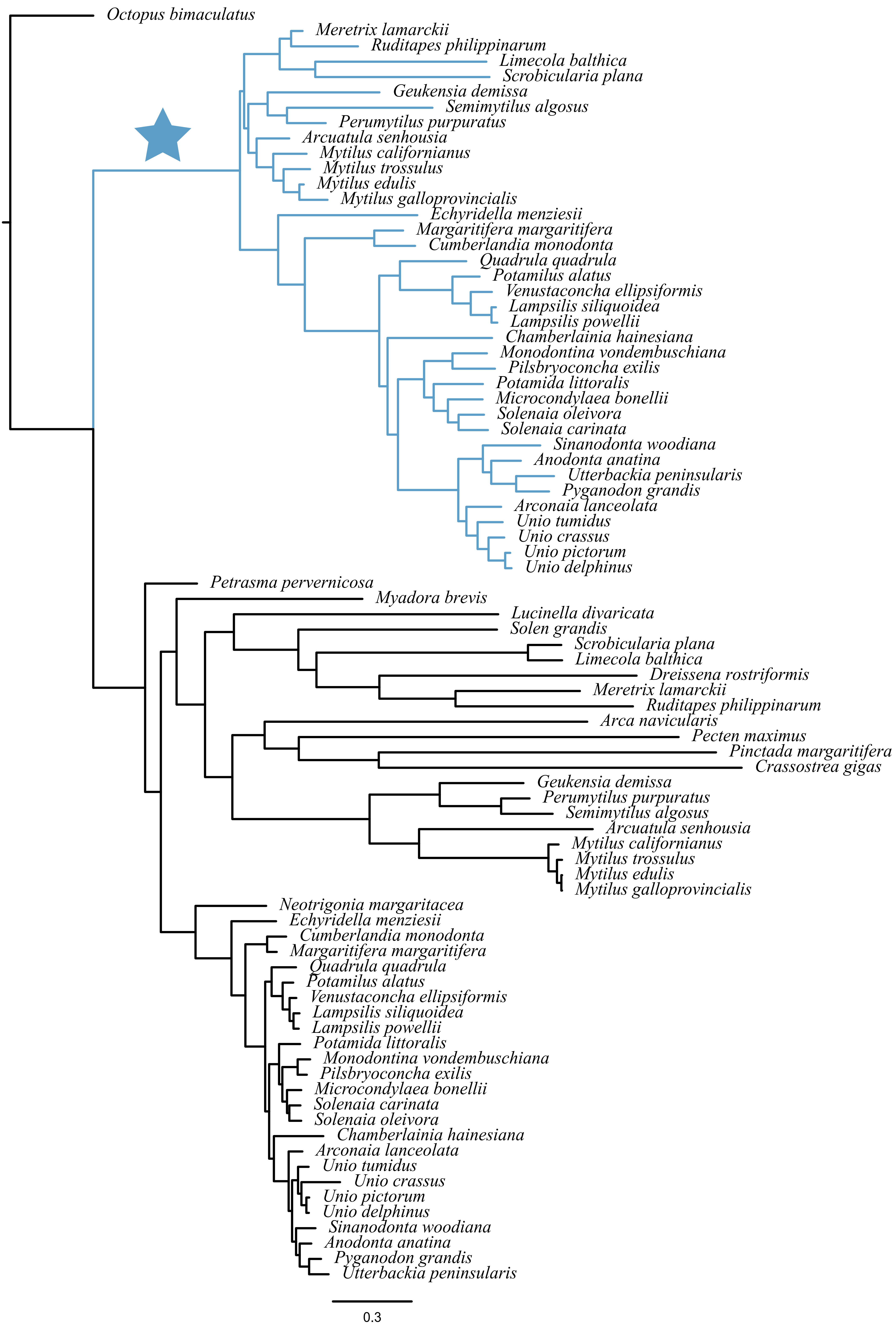

### FigS2_Mytilida_dated.pdf

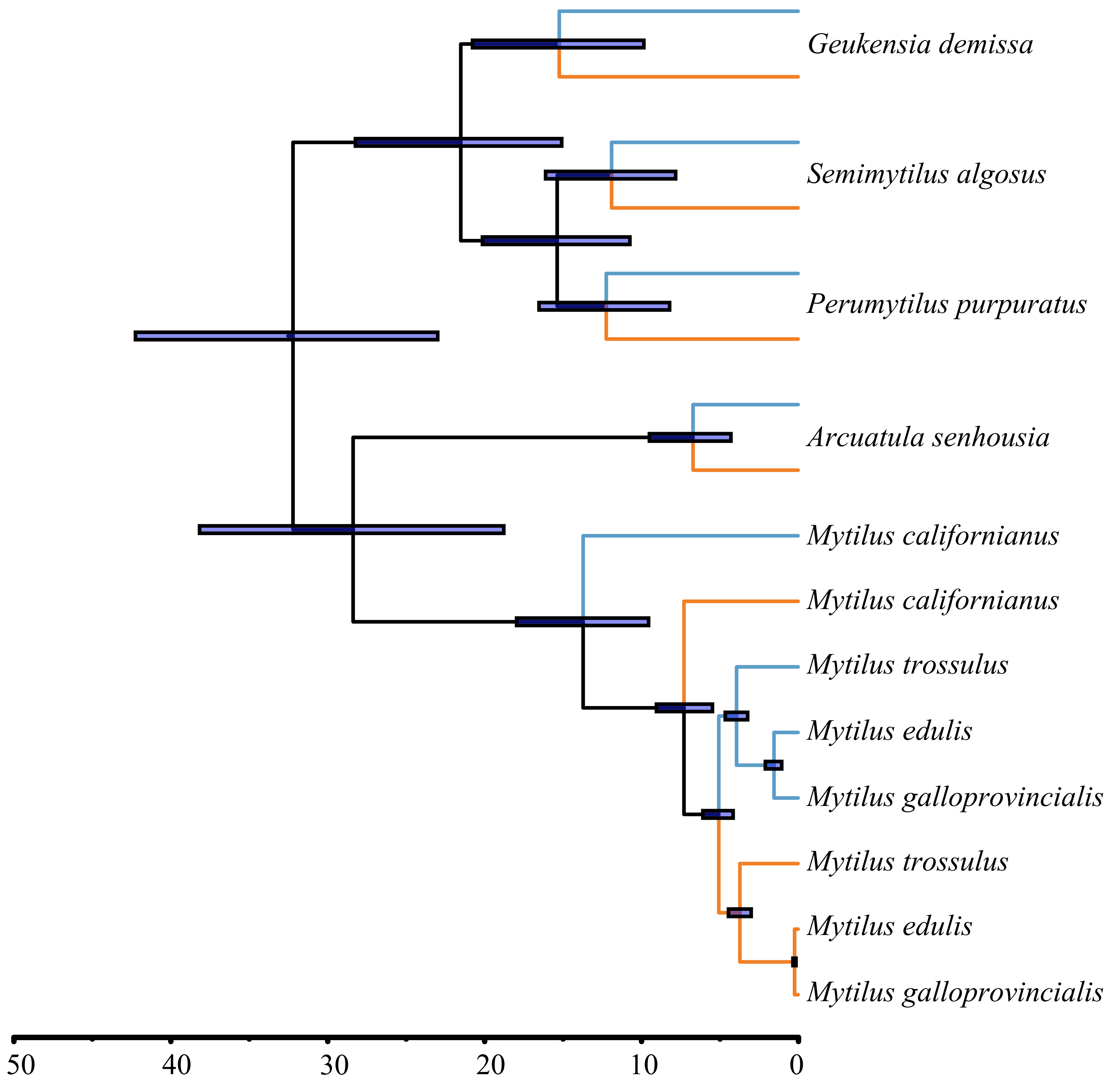

### FigS3_All_scf_plots.pdf

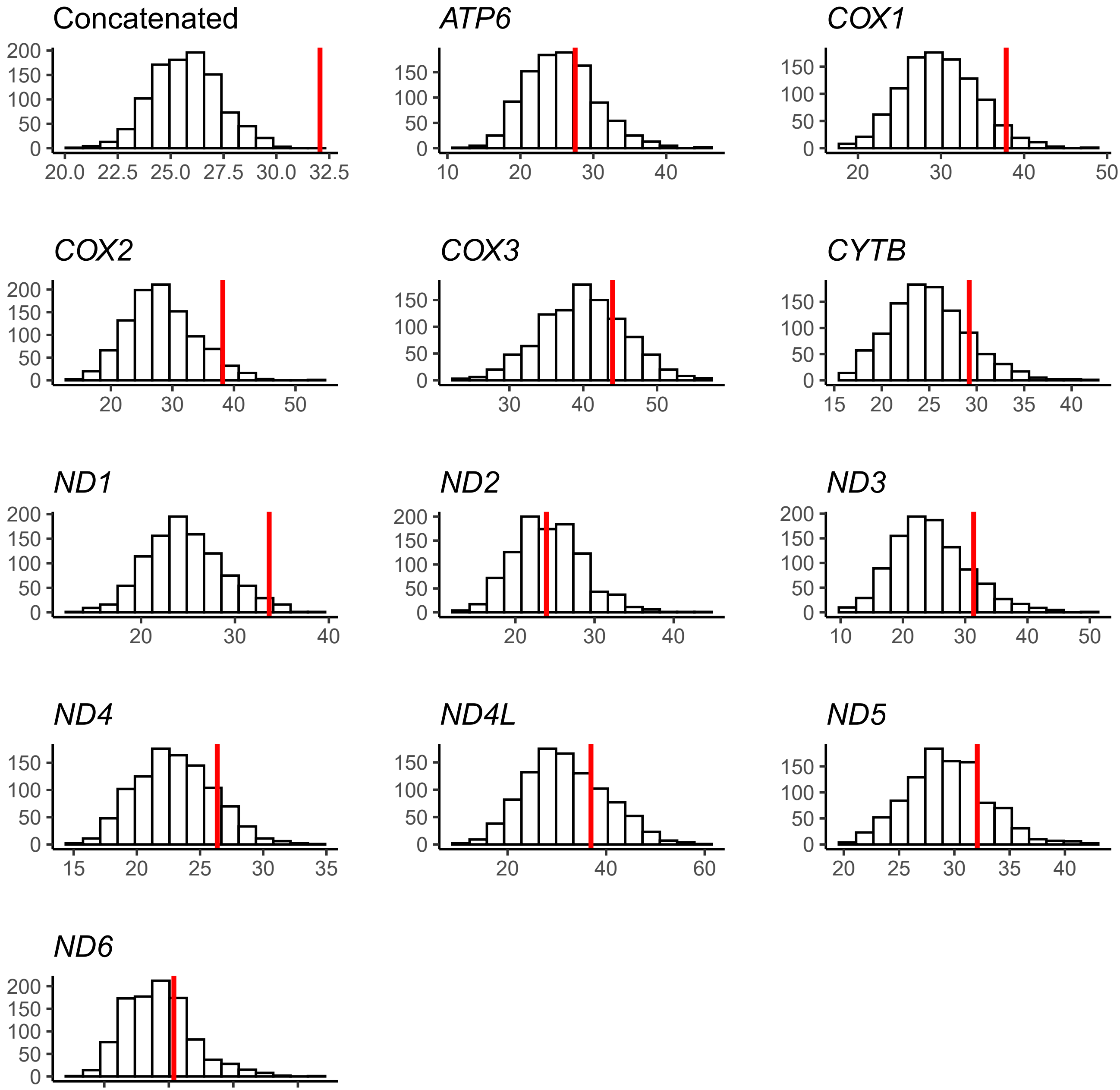
